# Placental pathology, circadian biology, and pathogenesis of spontaneous preterm birth: a pilot study of human placental gene expression profiling using a targeted HTG transcriptome panel

**DOI:** 10.64898/2026.06.23.734020

**Authors:** Guoli Zhou, Hanne Hoffmann, Hidemi Yamamoto, Kathryn Woods, Melanie Adkins, Robert Barbieri, Raina Nakova Fichorova

**Affiliations:** Center for Statistical Training & Consulting (CSTAT), Michigan State University, East Lansing, MI 48824; Department of Animal Science, Michigan State University, East Lansing, Ml 48824; Laboratory of Genital Tract Biology, Department of Obstetrics, Gynecology and Reproductive Biology, Brigham and Women’s Hospital, Harvard Medical School, Boston, MA 02115; Department of Obstetrics, Gynecology and Reproductive Biology, Brigham and Women’s Hospital, Harvard Medical School, Boston, MA 02115; Department of Epidemiology and Biostatistics, Michigan State University, East Lansing, Ml 48824

**Author notes:** **Co-Corresponding Authors:** Dr. Guoli Zhou, MPH, MD, PhD. Center for Statistical Training & Consulting (CSTAT), Michigan State University, East Lansing, MI 48824., Dr. Raina Nakova Fichorova, MD, PhD Brigham and Women’s Hospital, Harvard Medical School, Boston, MA 02115.

## Abstract

**BACKGROUND:** Spontaneous preterm birth (sPTB) remains the foremost cause of neonatal morbidity and mortality worldwide. Although histologic chorioamnionitis (HCA) and placental vascular abnormalities are frequently observed in sPTB, the molecular cascades linking these lesions to labor initiation remain poorly understood. Emerging evidence implicates circadian dysregulation and trophoblast dysfunction as additional drivers of sPTB.

**OBJECTIVE:** This study aims to map placental pathology to distinct transcriptomic functional signatures that may precipitate sPTB, delineate the contribution of circadian regulation - both core-clock genes and circadian transcription-factor target sets (TFTs) - to sPTB, and identify placental cell-type-enriched and developmental pathway signatures that differ between sPTB and term deliveries.

**STUDY DESIGN:** We performed bulk RNA sequencing on 32 formalin fixed, paraffin embedded placental specimens from 12 selected women (9 sPTB and 3 Term) in the POUCH Study cohort. Samples were selected for white ethnicity, maternal age 23-33 years, and parity 1-4 to reduce heterogeneity within groups. An extraction-free HTG transcriptome panel assayed 19,398 protein-coding genes. Log2-fold changes of all genes were computed with limma adjusted for maternal age, gestational age, parity, placental region, placental pathology, and POUCHID (a clustering variable) for sPTB vs. Term and HCA/vascular lesion vs. no pathology (no placental pathology adjustment). Gene-set enrichment used 50 Hallmark sets (MSigDB) plus curated placental circadian, circadian TFT, cell-type, and developmental pathways or gene sets.

**RESULTS:** sPTB placentas displayed a global suppression of metabolic, secretory, and immune pathways (e.g., protein secretion, oxidative phosphorylation, Interferon responses, Complement, ROS, MYC Targets, TGF β, mTORC1, and Coagulation) while KRAS Signaling Down and EMT were up-regulated. HCA-enriched sets (TNFα/NF-κB, ROS, KRAS Up, IL-2/STAT5, Hypoxia, Interferon-γ) were up-regulated, with EMT and Notch remaining down. Vascular abnormalities alone showed up-regulation of 12 Hallmark sets - including TGF-β, TNFα/NF-κB, ROS, pancreatic β-cell stress, Hypoxia, Oxidative Phosphorylation, EMT, and mTORC1 - while Notch was down-regulated. When HCA co-exists with vascular abnormalities, the Hallmark profile becomes more inflammatory highlighting a synergistic exacerbation of innate immunity, oxidative stress, and programmed cell death with the 12 up-regulated sets (Complement, Interferon α/γ, TNFα, ROS, Apoptosis, and Heme Metabolism). The exclusive downregulation of DNA Repair suggests compromised genomic integrity. Circadian gene-sets analysis revealed an up-regulated Regulation of Circadian Sleep Wake Cycle in sPTB but down-regulation of core clock pathway and suppressed circadian TF targets. Cell-type enrichment reveals increased trophoblast giant cells and IGFBP1-DKK1 positive fetal cells, with marked suppression of extravillous trophoblasts, syncytiotrophoblasts, villous cytotrophoblasts, and fetal myeloid cells. Placental developmental pathways were downregulated, indicating arrested trophoblast maturation.

**CONCLUSION:** Our pilot analysis demonstrates sPTB placentas exhibit a global suppression of metabolic, secretory, and immune-modulatory programs and maladaptive trophoblast remodeling, whereas HCA and vascular abnormalities drove distinct inflammatory or hypoxic signatures. The shared and opposing Hallmark pathways across phenotypes highlight distinct yet overlapping pathogenic mechanisms. Dysregulated circadian pathways, consistent downregulated transcription factor target gene sets, and trophoblast-specific signatures implicate circadian misalignment and impaired placental maturation as key contributors to preterm parturition. These findings provide a mechanistic atlas linking placental pathology to sPTB and highlight potential targets for chronotherapeutic and cell-type-specific interventions.

**AJOG at a Glance:** *Why was this study conducted?:* Spontaneous preterm birth remains a leading cause of neonatal morbidity. Histopathologic lesions of the placenta, particularly chorioamnionitis and vascular abnormalities, are common in preterm deliveries, yet the underlying molecular pathways are poorly understood. We sought to integrate functioning pathway profiles of placental histology, circadian biology, and cell types to identify mechanistic drivers of sPTB.

*Key findings:*

- sPTB placentas showed widespread down-regulation of oxidative phosphorylation, mTORC1, hypoxia, interferon, and TNFα/NF-κB pathways.
- HCA placentas up-regulated the same pathways (except androgen response), revealing a reciprocal inflammatory–hypoxic signature.
- Vascular abnormalities displayed a distinct mix of up- and down-regulated pathways, suggesting divergent reparative responses.
- Placentas with co-existing HCA and vascular abnormalities enriched more inflammatory Hallmark pathways: the 12 up-regulated sets (Complement, Interferon α/γ, TNFα, ROS, Apoptosis, and Heme Metabolism) highlight a synergistic exacerbation of innate immunity, oxidative stress, and programmed cell death and the exclusive down-regulation of DNA Repair suggests compromised genomic integrity, which can contribute to premature placental senescence and preterm labor.
- Circadian clock and multiple transcription-factor targets were enriched in sPTB, and trophoblast-specific signatures (giant, extravillous, syncytiotrophoblast) were prominent.

*What does this add to what is known?:* The study demonstrates a clear dichotomy between inflammatory and hypoxic molecular programs in sPTB and HCA, identifies circadian dysregulation as a potential contributor, and highlights trophoblast subpopulations as key players. These insights open avenues for targeted biomarkers and chronotherapy in preterm birth prevention.

## Introduction

Preterm birth (PTB), defined as delivery before 37 weeks of gestation, remains the leading cause of neonatal morbidity and mortality worldwide. PTB is a multifactorial and heterogeneous condition, clinically classified into spontaneous (sPTB, often accompanied by preterm premature rupture of membranes [PPROM]) and non-spontaneous or medically indicated PTB ^1–3^. In the United States, ∼2/3 of preterm births are spontaneous ^4^ with greater risk for women living in poverty and for African-American women ^5, 6^. Despite advances in obstetric care, the global PTB rate has plateaued at ∼10 %, ^7^ underscoring the urgent need to elucidate the underlying biological mechanisms that drive PTB.

A growing body of evidence implicates the placenta as a pivotal mediator of PTB, with histopathologic lesions - most notably histologic chorioamnionitis (HCA) and placental vascular abnormalities - constituting the most frequently observed pathological substrates in preterm placentas. ^8, 9^ Yet the molecular cascades linking these lesions to the initiation of labor remain incompletely understood. Molecular mechanisms contributing to sPTB in the absence of HCA and vascular abnormalities are even less studied.

The placenta is a dynamic organ that orchestrates maternal-fetal communication through endocrine, immune, and vascular functions. Placental lesions can arise from intrauterine infection, hypoxia, or maladaptive angiogenesis, each perturbing placental homeostasis and triggering inflammatory or hypoxic signaling pathways that culminate in preterm parturition. ^10–12^ HCA, characterized by neutrophilic infiltration of the chorionic plate and amniotic membranes, activates toll-like receptor (TLR) signaling, leading to the production of pro-inflammatory cytokines (IL-6, IL-1β) and matrix-remodeling enzymes that weaken cervical tissue and stimulate uterine contractility. ^13, 14^ Conversely, vascular lesions such as maternal vascular obstructive lesions (MV-O) or fetal vascular obstructive lesions (FV-O) compromise placental perfusion, inducing oxidative stress and hypoxia-inducible factor (HIF) pathways that have been linked to labor initiation. ^15, 16^ Moreover, oxidative stress and inflammation have been associated with systemic circadian dysregulation ^17^ and with disruption of placental clock gene expression, which in turn is linked to aberrant cytokine production and impaired angiogenesis. ^18, 19^ These processes are central to HCA and vascular injury in the pathophysiology of PTB. Circadian disruption, whether due to shift work, jet lag, or chronic jet-lag-like patterns, malnutrition or stress, ^20–22^ augments the risk of PTB, ^23, 24^ yet molecular mechanisms are poorly understood. Maternal circadian rhythms are known to influence placental clock machinery, ^25^ which comprises core transcription factors such as CLOCK, BMAL1, PER1/2, and CRY1/2 and regulates rhythmic expression of genes involved in hormone synthesis, nutrient transport, and immune tolerance regulating reproductive events, including ovulation, implantation, uterine contractions, and labor. ^18, 26–28^ However, their placental expression and role in PTB pathways have been sparsely studied.

Beyond global placental function, the roles of specific cellular compartments that may differentially contribute to PTB in the presence or absence of HCA or vascular injury are yet to be defined. Trophoblast subpopulations—cytotrophoblasts, syncytiotrophoblasts, extravillous trophoblasts (EVTs), and trophoblast-derived endothelial cells—undergo tightly regulated differentiation pathways that determine placental invasion, vascular remodeling, and immune tolerance. ^29–31^ Dysregulation of Notch, Wnt/β-catenin, and TGF-β signaling in the trophoblast has been implicated in impaired villous development and inadequate vascularization, features frequently observed in preterm placentas. ^32, 33^ Moreover, immune cell populations within the intervillous space—macrophages, neutrophils, and natural killer (NK) cells—play a dual role, mediating both protective immune surveillance and potential inflammatory triggers of labor when dysregulate. ^34–36^

We hypothesized that, by interrogating placental cell-type signatures and developmental pathways with the curated gene-set enrichment analyses, we could pinpoint the specific cellular and circadian contexts in which sPTB-associated transcriptional changes arise. To test this hypothesis, we leveraged a robust, global transcriptome analysis along with the detailed histopathologic diagnosis and clinical metadata of the well-characterized Pregnancy Outcomes and Community Health (POUCH) Study cohort. ^37^ Our objectives are threefold: (i) to map the placental histopathologic lesions to distinct transcriptomic functional gene sets or pathways that may precipitate sPTB; (ii) to delineate the contribution of circadian regulation—both core clock genes and circadian transcription-factor target sets—to sPTB vs. Term; and (iii) to identify placental cell-type-enriched and developmental pathway signatures that differ between sPTB and term deliveries. By integrating Hallmark gene-set enrichment, circadian pathways, circadian transcription-factor targeting (TFT) gene sets, and placental cell types and developmental pathway analyses within a rigorous statistical framework that accounts for maternal age, parity, gestational age, placental pathology, and intra-placental heterogeneity, we aim to generate a mechanistic atlas that links the placental histopathologic milieu to the molecular cascades of preterm labor.

## Methods

### Study Population and Sampling of Formalin-Fixed Paraffin-Embedded (FFPE) Placental Tissues

The POUCH Study cohort included 3,019 pregnant women enrolled between their 16th and 27th week of pregnancy from 52 clinics in Michigan (August 1998–June 2004). ^38^ Gestational age at delivery was calculated using the last menstrual period (LMP); ultrasound data (< 25 weeks) were used if the LMP and ultrasound estimates differed by > 2 weeks. ^39^

Michigan State University’s (MSU) IRB approved the POUCH Study and HIPAA Security Guidelines were applied for the deidentification of all data and the use of placental FFPE samples for transcriptomics in the current study. The Brigham and Women’s Hospital IRB approved the nested study using deidentified placental tissues in Fichorova’s Laboratory for the transcriptome analysis approved by the MSU protocol.

For the purposes of this nested transcriptome analysis we excluded PTB due to medical indications or premature rapture of the membranes and focused on sPTB defined as delivery before 37 weeks of gestation initiated by spontaneous labor (onset of regular contractions resulting in cervical dilation of at least 2 cm). ^39^ The study design illustrated in Figure 1 included 12 pregnancies (9 sPTB and 3 term deliveries) and 32 placental tissues representing no pathology, HCA and/or vascular abnormalities. The sample characteristics are shown in Table 1. To minimize the number of potential confounders the nested study was limited to white race and a narrow maternal age range at first prenatal visit (23.2-32.5) (Table 1). Placental regions sampled included the central area (with or without lesions), the cord-insertion area, and the marginal area with lesions. Placental pathology was recorded as HCA, vascular lesion, or no pathology. HCA was defined as ‘Yes’ = severe (polymorphonuclear leukocyte inflammatory pattern in chorionic plate and/or extraplacental membranes, plus karyorrhexis or necrotizing inflammation) and ‘No’ = mild (≥ 10 polymorphonuclear leukocytes in a high-power field) or none, due to no association between mild HCA and PTB. ^39, 40^ Vascular abnormality was defined as ‘Yes’ = presence of at least one of five placental vascular constructs - maternal or fetal obstructive lesions (MV-O or FV-O), maternal bleeding/vessel integrity (MV-I), lack of physiologic conversion of maternal spiral arteries (MV-D), and fetal bleeding/vessel integrity (FV-I) (Kelly et al., 2009) and ‘No’ otherwise.

**Figure 1.**
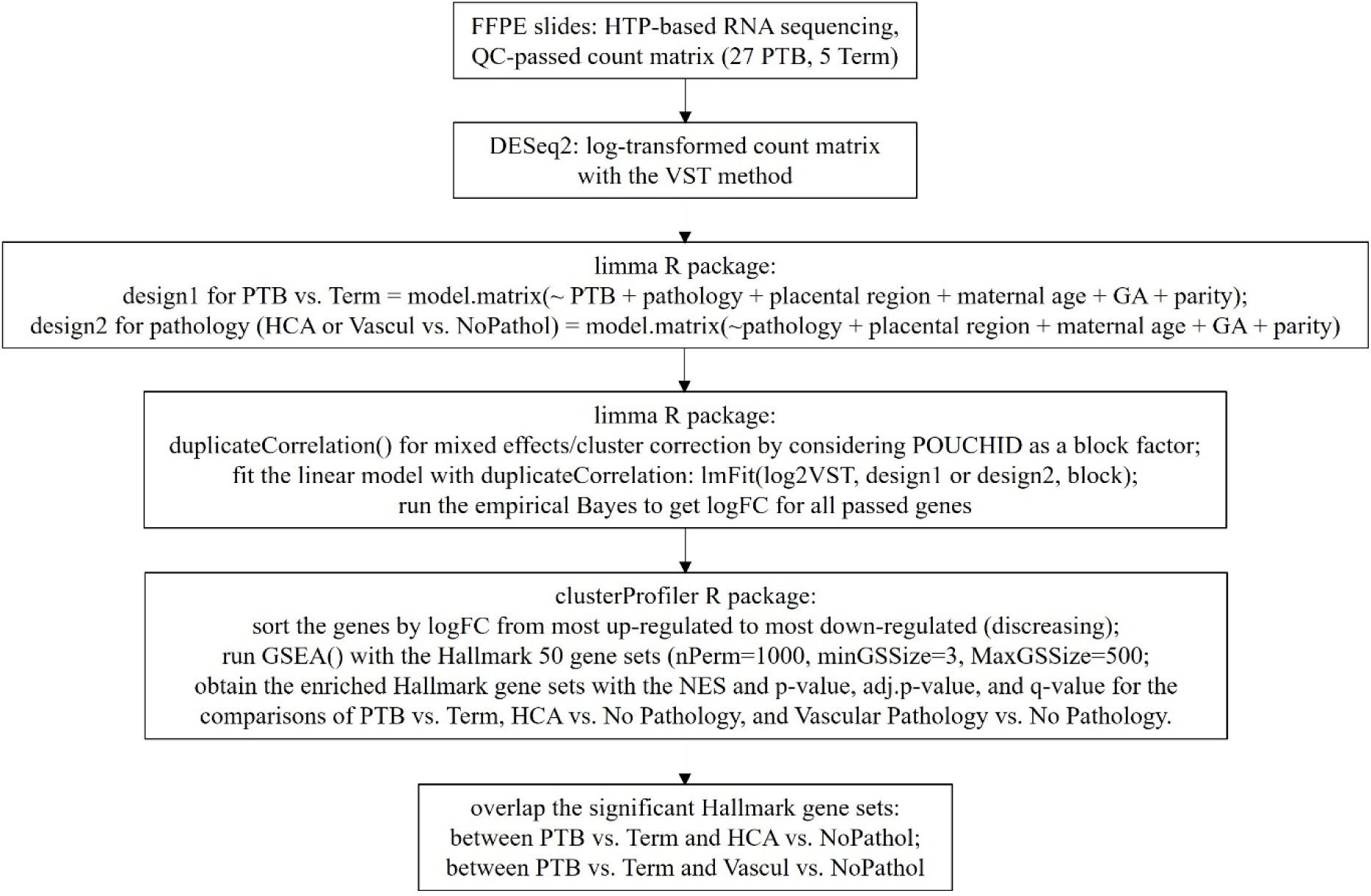
Flowchart illustrates study design and bioinformatics pipeline for analysis of placental gene expression. PTB=preterm birth; HCA=histologic chorioamnionitis; GA=gestational age

**Table 1.** Sample characteristics (N=32).

| sPTB vs. Term |  |  |  |  |
| --- | --- | --- | --- | --- |
|  |  | 0=Term (N=5) | 1=sPTB (N=27) | Overall (N=32) |
| Maternal age at delivery |  |  |  |  |
|  | Mean (SD) | 26.0 (2.66) | 28.6 (2.40) | 28.2 (2.58) |
|  | Median [Min, Max] | 27.4 [23.2, 28.9] | 28.3 [24.4, 32.5] | 27.9 [23.2, 32.5] |
| Final GA at delivery (weeks) |  |  |  |  |
|  | Mean (SD) | 39.1 (1.36) | 34.0 (0.862) | 34.8 (2.09) |
|  | Median [Min, Max] | 38.9 [38.1, 41.4] | 34.4 [32.7, 35.0] | 34.4 [32.7, 41.4] |
| Parity |  |  |  |  |
|  | 1 | 0 (0%) | 10 (37.0%) | 10 (31.3%) |
|  | 2 | 5 (100%) | 10 (37.0%) | 15 (46.9%) |
|  | 3 | 0 (0%) | 4 (14.8%) | 4 (12.5%) |
|  | 4 | 0 (0%) | 3 (11.1%) | 3 (9.4%) |
| Pathology |  |  |  |  |
|  | None | 5 (100%) | 8 (29.6%) | 13 (40.6%) |
|  | HCA only | 0 (0%) | 2 (7.4%) | 2 (6.3%) |
|  | Vascular only | 0 (0%) | 14 (51.9%) | 14 (43.8%) |
|  | Both | 0 (0%) | 3 (11.1%) | 3 (9.4%) |
| Placental Region |  |  |  |  |
|  | central disc with lesions | 1 (20.0%) | 6 (22.2%) | 7 (21.9%) |
|  | central disc normal | 2 (40.0%) | 9 (33.3%) | 11 (34.4%) |
|  | disc at cord insertion | 1 (20.0%) | 7 (25.9%) | 8 (25.0%) |
|  | marginal with lesions | 1 (20.0%) | 5 (18.5%) | 6 (18.8%) |

### Evaluation of RNA Quality

The DV200 metric is a precise measure of RNA fragmentation (particularly with the FFPE samples), defined as the percentage of total RNA fragments > 200 nucleotides in length (Illumina Technical Note. 2014). To determine the quality of total RNAs from the archived FFPE tissues, we randomly selected 3 POUCH term placental FFPE blocks and then 3 sections (10-μm thickness/section) per tube (1.5-ml, RNase-free) from the selected blocks were cut using the microtome. The total RNAs were extracted from the sections by the NucleoSpin total RNA FFPE XS kit (TaKaRa Bio), according to the manufacturer’s manual. The High Sensitivity RNA ScreenTape with the TapeStation Analysis Software 5.1 was used to assess the DV200 metric (Agilent Technologies, Inc. 2023). The results were summarized as average DV200 with a standard deviation (SD) as well as the minimum/maximum DV200.

### Global mRNA Transcriptome Assay

Global mRNA transcriptomes of >19,000 human genes were profiled in Dr. Fichorova’s Laboratory of Genital Tract Biology using the High Throughput Genomics Inc (HTG) nuclease protection (HTP) assay, which has been extensively validated for human FFPE tissue. ^41–45^ The HTP assay obviates RNA extraction; a small scraped FFPE section amplifies the entire transcriptome. It contains probes for 19,398 genes plus 218 control probes (4 positive, 100 negative, 22 genomic DNA, 92 ERCC). For each placental block, consecutive 10-µm sections were mounted on charged slides: the first was H&E-stained and scanned on a Zeiss Axioscan 7 at 10× (Digital Slide Scanner). The .czi files were converted to .tiff with QuPath ^46^ and imported into ImageScope to estimate the area of the adjacent unstained section. That section was stripped from the slide, embedded in HTG lysis buffer to achieve 22 mm²/50 µl, centrifuged (1 min), heated to 95 °C (20 min), cooled 10 min, treated with proteinase K (5 %) and incubated at 50 °C for 3 h on an orbital shaker. After freezing at –80 °C, samples were processed on the EdgeSeq platform: protection probes hybridized to mRNA, amplified, tagged, and quantified by qPCR. Libraries (Kapa Library Quantification Kit) were sequenced on an Illumina NextSeq 500. Raw data were parsed from *.fastq files with HTG EdgeSeq Parser and QC’d in EdgeSeq Reveal, using thresholds ≥10 % unique mapping and ≥30 % on-target reads.

### Generation of log2-fold changes of all detectable genes via differential expression analysis

The QC-passed raw count matrix was subjected to a variance-stabilizing transformation (VST) and log2-transformation with the DESeq2 package. ^47^ Log2-fold changes (logFCs) of all genes were estimated with the limma package. ^48^ Studies have shown that both placental pathology and placental sampling region are potential confounders in the relationship between preterm birth and placental gene expression/pathways. ^31, 49–52^ Meanwhile, evidence also indicated that placental gene expression is gestational age (GA)-dependent.^51^ Thus, in order to isolate pathologenic “preterm-vs-term delivery” effects from the normal GA-dependent developmental program, the GA was also considered a potential confounder. Two design matrices were specified: (1) PTB vs. Term, incorporating confounders (pathology, placental region, maternal age, gestational age, parity); (2) pathology comparison (HCA or vascular abnormality vs. no pathology), with the covariates (placental region, maternal age, gestational age, parity). Because samples from the same donor (POUCHID) were not independent, the ‘duplicateCorrelation’ function from the limma package was applied to estimate and adjust for within-donor correlation, treating POUCHID as a block factor. ^48, 53^ The linear model was fitted with lmFit, followed by empirical Bayes moderation (eBayes) to yield adjusted log2-fold changes (logFCs) and moderated t-statistics for all genes for four comparisons (sPTB vs. Term, HCA vs. no pathology, vascular lesion vs. no pathology, both HCA and vascular lesion vs. no pathology), respectively (Table S1-S4).

### Hallmark Gene-Set Enrichment Analysis

Among the gene-set collections in MSigDB, ^54, 55^ the Hallmark 50 sets offer distinct advantages: they reduce noise and redundancy, are curated from multiple founder sets to convey specific biological states with coherent expression, are interpretable (only 50 sets for detailed review), and are reproducible with well-documented definitions. ^56^ In R, the msigdbr function (species = “Homo sapiens”, category = “H”), ^57^ was used to retrieve these sets, producing a data frame (TERM2GENE) with term IDs and mapped genes. All 19,398 genes were ranked by logFCs (adjusted for maternal age, gestational age at visit, parity, placental pathology, placental region, and POUCHID as a block factor) and fed into the GSEA function in clusterProfilerpackage. ^58^ GSEA utilizes the full ranked list rather than arbitrary DEGs, enabling detection of subtle coordinated changes across pathways, avoiding missed biologically relevant pathways, and preventing bias from excluding genes with small yet meaningful changes. Normalized enrichment scores (NES), nominal p-values, FDR-adjusted p-values, and q-values were extracted for all 50 Hallmark sets per comparison. Significant sets were defined as p < 0.05 and adjusted p < 0.10, then mapped between PTB-enriched, HCA-enriched, vascular lesion-enriched, and both HCA/vascular lesion-enriched comparisons to link placental pathology to sPTB.

### Curated Circadian, TFT, Cell-Type, and Developmental Gene Sets

Restricting GSEA to the four curated placenta-centric modules improves tissue specificity, biological resolution, and reduces statistical noise—essential for placenta-related disease research—whereas generic MSigDB collections are broad, often irrelevant, and can mask true signals due to redundancy and inflated multiple-testing penalties.

Circadian pathways were curated by searching “circadian” on the MSigDB Browse Human Gene Sets page (https://www.gsea-msigdb.org/gsea/msigdb/human/genesets.jsp), yielding 21 pathway names across collections. Using the R msigdbr package, ^57^ the sets were merged and subjected to GSEA on the ranked logFCs of 19 398 genes comparing sPTB to Term, identifying sPTB-enriched circadian pathways.

We retrieved the C3 subcollections ‘TFT:TFT_LEGACY’ (506 sets) and ‘TFT:GTRD’ (610 sets), extracted unique TF symbols, and mapped them to the human circadian gene list (1,409 genes) from the CGDB database. ^59^ This yielded 67 circadian TF-target sets (TFTs), which were tested by GSEA on the same logFC ranking to find sPTB-enriched circadian TFTs.

Finally, placental cell-type signatures and developmental pathways were curated by searching the keywords “trophoblast,” “placenta,” and “placental” in the Gene Set Name field. This produced 16 placental cell-type gene sets and 23 placental developmental pathways. GSEA on the sPTB vs. Term logFCs highlighted enriched placental cell-type and developmental gene sets.

### Correlations of significant PTB-enriched Hallmark pathways with significant PTB-enriched circadian gene sets including circadian pathways and circadian TFTs

Variance stabilized log2(VST) matrix generated by the DESeq2 and MSigDB Hallmark gene sets/curated circadian gene sets (circadian pathways + circadian TFTs) were input into the GSVA package [1] ^60^ to obtain GSVA pathway activity score matrices (i.e., z-score matrices) at the individual sample level. The z-score matrices were further subset by the significant PTB-enriched pathways or gene sets followed by correlation analyses of significant PTB-enriched Hallmark pathways with significant PTB-enriched circadian gene sets (circadian pathways + circadian TFTs) using the rcorr function (type=’pearson’) in the Hmisc package. ^61^ Heatmaps of the correlation matrices were visualized by the pheatmap package.

All analyses were conducted in R (version 4.3.2).

## Results

### RNA Quality

The mean DV200 across three FFPE term placentas was 48.5 % (SD = 14.5 %; minimum 34.7 %, maximum 70.0 %) (data not shown). Evidence indicates that, by adjusting RNA input amounts, high-quality cDNA libraries can be prepared from FFPE RNA samples with DV200 ≥ 25 %. ^62–64^ Thus, our DV200 metric assessment indicates that the POUCH-archived FFPE tissue samples are highly suitable for bulk RNA-seq.

### Hallmark Pathways in sPTB

GSEA of sPTB vs. Term revealed 14 significant Hallmark sets (p < 0.05, adjusted p < 0.10). All but two were down-regulated: Protein Secretion, Oxidative Phosphorylation, Androgen Response, Interferon-α Response, Complement, Reactive Oxygen Species Pathway, Interferon-γ Response, MYC Targets V1, TGF-β Signaling, Allograft Rejection, mTORC1 Signaling, and Coagulation. Only KRAS Signaling Down and Epithelial-Mesenchymal Transition (EMT) were up-regulated (Table S5, Table 2).

**Table 2.** Summary of the GSEA with Hallmark Gene Sets from the Molecular Signatures Database (MSigDB) for the full data (27 Preterm Birth vs 5 Term).

| Hallmark Gene Set ID (sorted by NES*, lowest to highest) | PTB vs Term | HCA vs No Pathology | Vascular vs No Pathology | Both vs No Pathology |
| --- | --- | --- | --- | --- |
| HALLMARK_PROTEIN_SECRETION | Down |  | Up |  |
| HALLMARK_OXIDATIVE_PHOSPHORYLATION | Down |  | Up |  |
| HALLMARK_ANDROGEN_RESPONSE | Down |  |  |  |
| HALLMARK_INTERFERON_ALPHA_RESPONSE | Down |  |  | Up |
| HALLMARK_COMPLEMENT | Down |  |  | Up |
| HALLMARK_REACTIVE_OXYGEN_SPECIES_PATHWAY | Down | Up | Up | Up |
| HALLMARK_INTERFERON_GAMMA_RESPONSE | Down | Up |  | Up |
| HALLMARK_MYC_TARGETS_V1 | Down |  |  |  |
| HALLMARK_TGF_BETA_SIGNALING | Down |  | Up |  |
| HALLMARK_ALLOGRAFT_REJECTION | Down |  | Up | Up |
| HALLMARK_MTORC1_SIGNALING | Down |  | Up |  |
| HALLMARK_COAGULATION | Down |  |  |  |
| HALLMARK_KRAS_SIGNALING_DN | Up | Up |  |  |
| HALLMARK_EPITHELIAL_MESENCHYMAL_TRANSITION | Up | Down | Up | Up |
| HALLMARK_TNFA_SIGNALING_VIA_NFKB |  | Up | Up | Up |
| HALLMARK_PEROXISOME |  | Up |  | Up |
| HALLMARK_UV_RESPONSE_UP |  | Up | Up | Up |
| HALLMARK_IL2_STAT5_SIGNALING |  | Up |  |  |
| HALLMARK_HYPOXIA |  | Up | Up | Up |
| HALLMARK_NOTCH_SIGNALING |  | Down | Down |  |
| HALLMARK_PANCREAS_BETA_CELLS |  |  | Up |  |
| HALLMARK_UV_RESPONSE_DN |  |  | Up |  |
| HALLMARK_APOPTOSIS |  |  |  | Up |
| HALLMARK_HEME_METABOLISM |  |  |  | Up |
| HALLMARK_DNA_REPAIR |  |  |  | Down |

### Hallmark Pathways in HCA and Vascular Abnormality

Stratifying PTB by HCA revealed that almost all HCA-enriched Hallmark pathways were up-regulated (except for Epithelial Mesenchymal Transition and Notch Signaling pathways) (Table S6, Table 2). Up-regulated sets included TNFA Signaling via NF-KB, Reactive Oxygen Species Pathway, Peroxisome, Kras Signaling Up, UV Response Up, Allograft Rejection, IL2/STAT5 Signaling, Hypoxia, and Interferon Gamma Response.

Placental vascular abnormalities (MV-O, FV-O, MV-I, FV-I, MV-D) were associated with significant enrichment of twelve up-regulated Hallmark gene sets—TGF Beta Signaling, TNFα Signaling Via NF-kB, Reactive Oxygen Species Pathway, Pancreas Beta Cells, Allograft Rejection, Protein Secretion, Hypoxia, Oxidative Phosphorylation, UV Response Dn, Epithelial Mesenchymal Transition, MTORC1 Signaling, and UV Response Up—and one down-regulated set—Notch Signaling (Table S7, Table 2).

The transcriptome of the placentas with both HCA and at least one of 5 vascular abnormalities in sPTB was enriched by 12 up-regulated Hallmark pathways (Interferon-α Response, Complement, Reactive Oxygen Species Pathway, Interferon-γ Response, Allograft Rejection, Epithelial Mesenchymal Transition, TNFA Signaling via NF-KB, Peroxisome, UV Response Up, Hypoxia, Apoptosis, and Heme Metabolism) and only DNA Repair pathway was down-regulated (Table S8, Table 2).

### Overlap of Hallmark Signatures

The analysis revealed shared enriched signatures between the two major placental histopathology types as well as their unique relationship to PTB in the absence of histopathology (Table 2). Four Hallmark gene sets were upregulated within the vascular abnormality-enriched and HCA-enriched signatures, shared among placentas that expressed either one or both of these histotypes. The shared dysregulation included Hypoxia, Reactive Oxygen Species, TNFα Signaling via NF-κB and UV Response UP, which are part of the heightened physiologic responses to DNA damage and stress. Among those, the Reactive Oxygen Species gene set was uniquely downregulated in the PTB-enriched gene signature suggesting impaired physiologic responses. In contrast, KRAS Signaling Down was up-regulated in both PTB and HCA and Epithelial-Mesenchymal Transition was upregulated in both PTB and in placentas with vascular abnormalities either alone or in combination with HCA. Of all 12 gene sets upregulated when both histotypes were present, five were enriched in the opposite direction (downregulated) in PTB in the absence of histopathology (Interferon-α or -γ Response, Complement, and ROS Pathway).

### Circadian Pathways and TFT Gene Sets in sPTB

sPTB-enriched circadian pathways included the up-regulated GO (BP) “Regulation of Circadian Sleep Wake Cycle” pathway and the down-regulated Reactome “Circadian Clock” pathway. The sPTB-enriched down-regulated circadian TFT gene sets comprised the target genes of the circadian transcription factors THRAP3, POU2AF1, ZNF146, and ARID5B (Table 3).

**Table 3.**
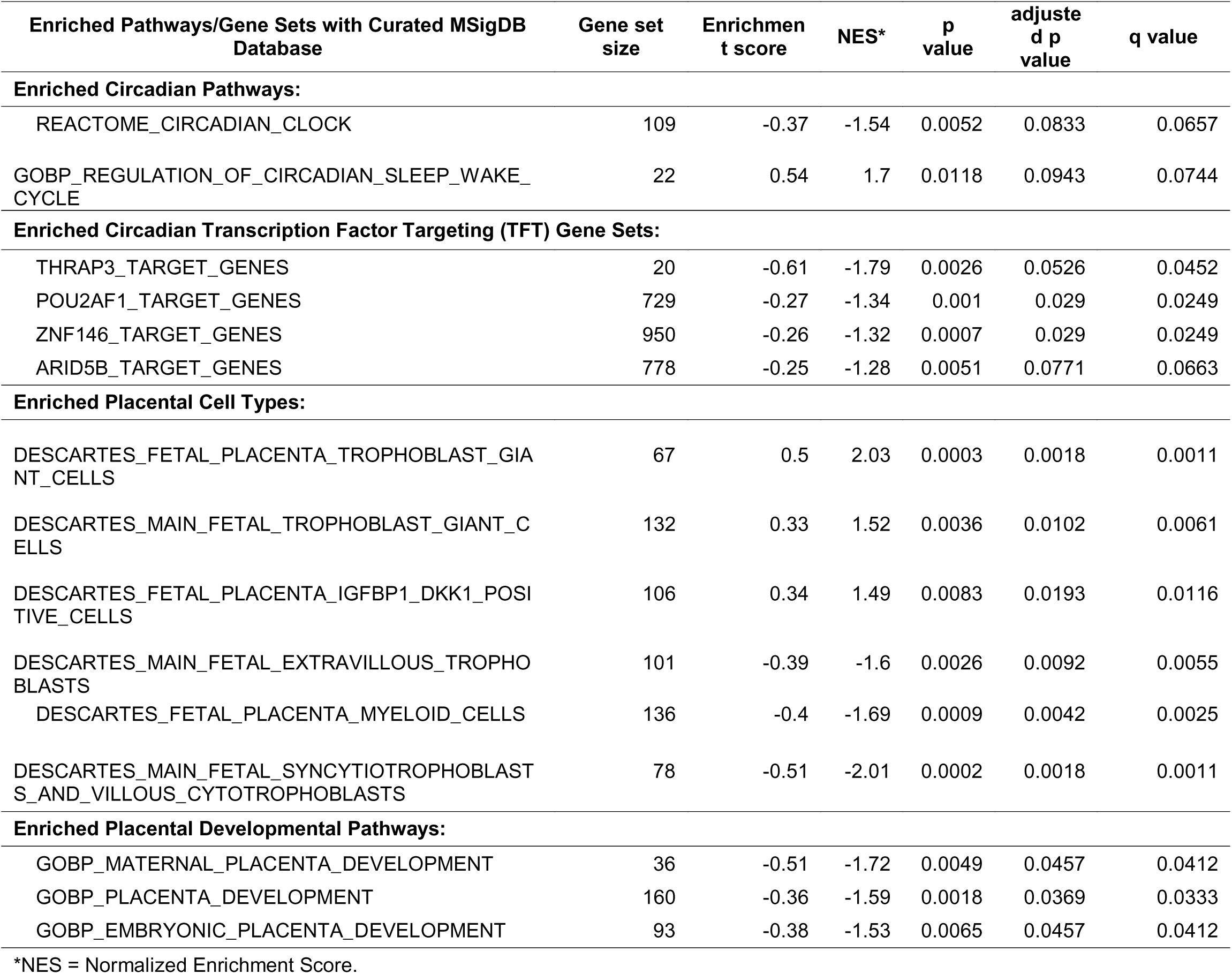
Enriched pathways with the curated MSigDB gene sets and the ranked log2FCs of all genes in PTB vs. Term adjusting for maternal age, parity, gestational age at the visiting, placental pathology, sampling regions of placenta, and POUCH ID (considered as a clustering variable).

### Placental Cell-Type and Developmental Signatures in sPTB

The cell-type enrichment analysis identified several placental populations that were over-represented in spontaneous preterm birth (sPTB) cases: up-regulated Trophoblast Giant Cells (main fetal and fetal subtypes) and Fetal Placenta IGFBP1 DKK1 Positive Cells as well as down-regulated Extravillous Trophoblasts (main fetal), Syncytiotrophoblasts and Villous Cytotrophoblasts (main fetal), and Fetal Placenta Myeloid Cells (Table 3). Gene-ontology enrichment highlighted the following down-regulated biological-process terms: Maternal Placenta Development, Placenta Development, and Embryonic Placenta Development (Table 3).

### Correlations Between Significant PTB-Enriched Hallmark Pathways and Significant PTB-Enriched Circadian Gene Sets

As shown in Figure 2 and Table S9, among 6 significant PTB-enriched circadian gene sets, the circadian clock pathway and circadian ARID5B/POU2AF1/ZNF146 TFTs were consistently, positively correlated with PTB-enriched protein secretion, oxidative phosphorylation, androgen response, complement, ROS, MYC v1, TGF beta pathway, and mTORC1 pathway (r=0.386 ∼ 0.659, p=0.0289 ∼ <0.0001) (Figure 2, Table S9), and negatively correlated with PTB-enriched KRAS pathway (r=-0.503 ∼ −0.546, p=0.0034 ∼ 0.0012) (Figure 2, Tables S9).

**Figure 2.**
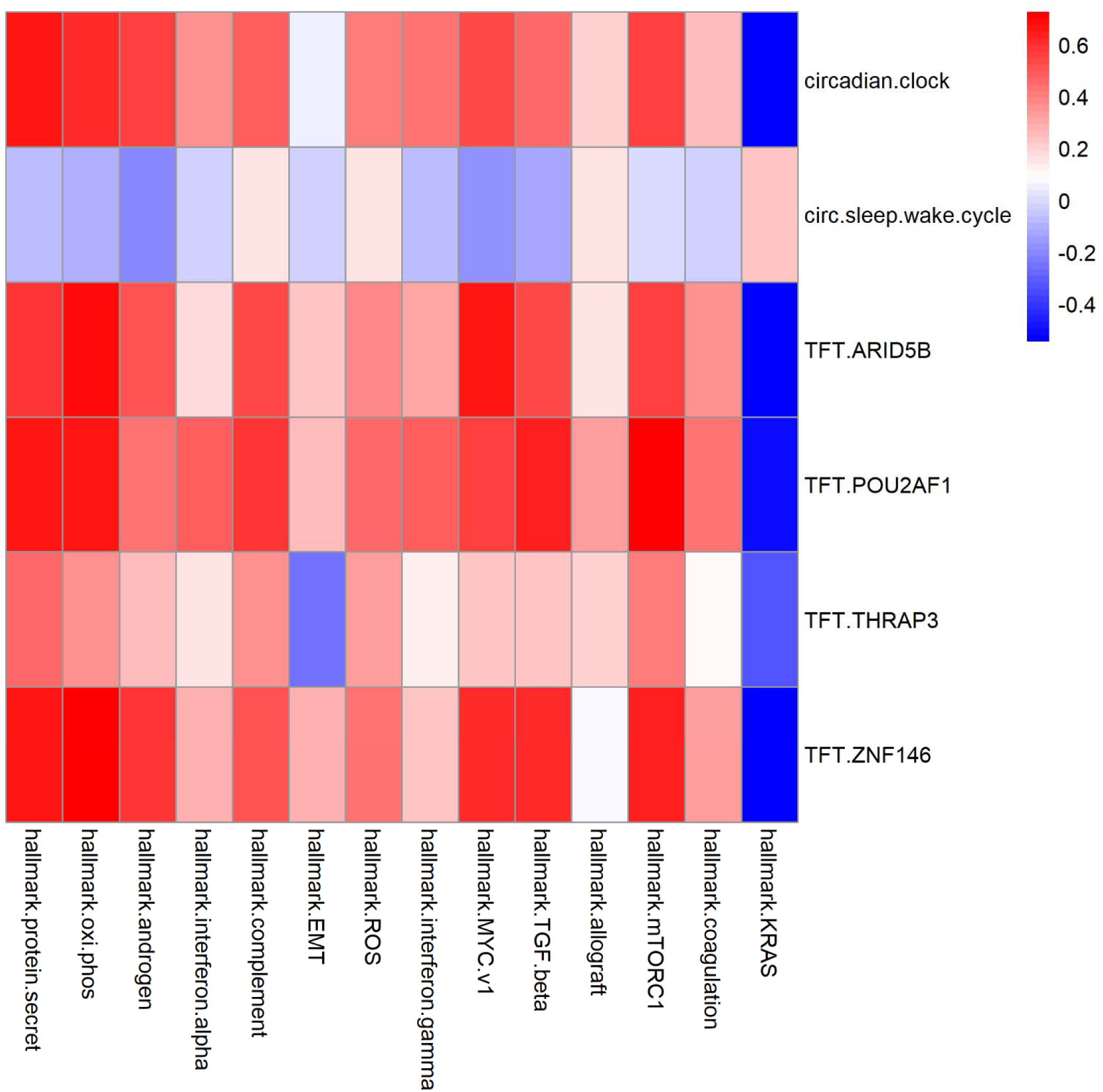
Heatmap illustrating correlations between significant preterm birth (PTB)-enriched hallmark pathways and significant PTB-enriched circadian gene sets

## Comment

### Principal Findings

Placental transcriptomes from sPTB compared to term birth in the setting of no significant histopathology show down-regulation of Hallmark pathways (Protein Secretion, Oxidative Phosphorylation, Interferon α and γ Responses, Complement, ROS, MYC Targets, TGF β, mTORC1, and Coagulation) while KRAS Signaling Down and EMT are up-regulated. Stratifying sPTB by HCA inverts the signature: most HCA-enriched sets (TNFα/NF-κB, ROS, KRAS Up, IL-2/STAT5, Hypoxia, Interferon-γ) become up-regulated, except EMT and Notch pathways. These data suggest the PTB placental gene dysregulation includes pathways driven by inflammation and vascular pathology but also allostasis occurring in the absence of histopathology, possibly independently driven by systemic maternal-fetal exposures. Vascular abnormalities exhibit an up-regulation of 12 Hallmark sets, including TGF β, TNFα/NF κB, ROS, pancreatic β cell stress, Protein Secretion, Hypoxia, Oxidative Phosphorylation, UV Down, EMT, and mTORC1; Notch is down. Overlap analysis shows KRAS Signaling Down is up-regulated in both sPTB and HCA, whereas ROS and Interferon-γ are oppositely regulated, and EMT is consistently up in HCA and vascular abnormalities but down in sPTB. When HCA co-exists with vascular abnormalities, the Hallmark profile becomes more inflammatory - 12 up-regulated sets (including Complement, Interferon α/γ, TNFα, ROS, Apoptosis, and Heme Metabolism) and one down-regulated DNA Repair pathway. Circadian gene-sets analysis links sPTB to an up-regulated “Regulation of Circadian Sleep Wake Cycle” term and a down-regulated core clock pathway, with suppressed circadian TF targets. Cell type enrichment reveals increased trophoblast giant cells and IGFBP1-DKK1 positive fetal cells, suppression of extravillous trophoblasts, syncytiotrophoblasts, villous cytotrophoblasts, and fetal myeloid cells; ontology terms are down-regulated, indicating arrest. Further correlation analysis revealed that the circadian clock pathway and circadian ARID5B/POU2AF1/ZNF146 TFTs consistently and positively correlated with PTB-enriched protein secretion, oxidative phosphorylation, androgen response, complement, ROS, MYC v1, TGF beta pathway, and mTORC1 pathway, and negatively correlated with PTB-enriched KRAS pathway. These alterations drive sPTB lesions leading to early uterine activation and fetal compromise, suggesting metabolic insufficiency, mitochondrial dysfunction, circadian disruption, and dysregulated immune/EMT signaling synergistically trigger premature labor.

## Results in context

### sPTB placentas exhibit a global suppression of metabolic, secretory, and immune-modulatory programs

The striking predominance of down-regulated Hallmark sets in sPTB—including Oxidative Phosphorylation, Protein Secretion, Complement, and both Interferon-α/γ responses—suggests that the preterm placenta operates under a state of energetic and functional compromise. Mitochondrial dysfunction has been repeatedly reported in sPTB placentas, with impaired electron transport chain activity, reduced ATP production, and increased reactive oxygen species (ROS). ^65–67^ Likewise, the down-regulation of the Protein Secretion pathway aligns with reduced syncytiotrophoblast secretory capacity, which is essential for hormone production and nutrient transport. ^26, 68^ The concurrent suppression of Complement and Coagulation pathways may reflect an anti-inflammatory or hypofunctional state that predisposes to early labor. ^69–71^ Interestingly, the androgen response pathway is also diminished, a finding that aligns with reports of altered steroidogenic signaling in sPTB placentas. ^65^ Together, these changes point to a placenta that is metabolically exhausted, less capable of sustaining normal trophoblast functions, and less prepared for the immune challenges that accompany parturition.

### Up-regulation of KRAS signaling and EMT in sPTB indicates maladaptive trophoblast remodeling

The only Hallmark pathways that were up-regulated in sPTB were KRAS Signaling Down and Epithelial-Mesenchymal Transition (EMT). KRAS activation has been implicated in trophoblast invasion and in the regulation of placental angiogenesis. ^33, 72, 73^ However, aberrant KRAS signaling can also drive proliferative and inflammatory phenotypes ^74^ that may contribute to premature labor. ^75–77^ EMT is a hallmark of trophoblast plasticity required for proper invasion of the uterine lining. ^78^ While EMT is normally tightly regulated, its ectopic or sustained activation can lead to pathological remodeling, excessive extracellular matrix degradation, and a pro-inflammatory milieu—all of which may precipitate sPTB. ^79–81^ The integration of these two up-regulated pathways with the down-regulated metabolic and secretory programs underscores a shift from a balanced, functionally competent placenta to one that is overly proliferative yet energetically compromised.

### HCA-enriched placentas show a classic infection-driven inflammatory signature

In contrast to the metabolic suppression seen in sPTB, HCA placentas displayed almost universal up-regulation of Hallmark pathways involved in innate immunity and oxidative stress. TNFA Signaling via NF-κB, Interferon-γ and Interferon-α responses, and the Complement system were all markedly activated, mirroring the host response to microbial invasion. ^56, 82^ Reactive Oxygen Species and Peroxisome pathways were also enriched, supporting the notion that HCA provokes oxidative damage and lipid peroxidation, ^83^ which can further amplify inflammation. ^9, 84–86^ The presence of IL2/STAT5 and UV Response Up sets indicates adaptive immune activation and DNA damage responses, respectively, consistent with the toll-like receptor (TLR)-mediated signaling cascades that have been described in chorioamnionitis. ^13, 87, 88^ These findings reinforce the concept that infection-induced inflammation is a primary driver of early labor in HCA, contrasting with the metabolic exhaustion model of sPTB.

### Placental vascular abnormalities are associated with a mixed oxidative-metabolic and remodeling signature

Vascular abnormalities such as maternal vascular malperfusion (MV-O) and fetal vascular malperfusion (FV-O) display a distinct constellation of up-regulated Hallmark sets. Enrichment of TGF-β Signaling, TNFα Signaling via NF-κB, and Hypoxia reflects the interplay between ischemia-reperfusion injury and profibrotic signaling ^89–93^ that may drive abnormal villous remodeling. ^91, 94^ The activation of UV Response Dn and UV Response Up alongside Pancreas Beta Cells and Allograft Rejection suggests a heightened DNA damage response and alloimmune reaction, ^95–98^ consistent with the endothelial injury seen in vascular abnormalities. ^99, 100^ Up-regulation of Epithelial-Mesenchymal Transition indicates a shift toward a mesenchymal phenotype in endothelial and trophoblast cells, promoting fibrosis and vascular remodeling. ^101–105^ The singular down-regulation of Notch Signaling further underscores the disruption of angiogenic pathways ^106^ that normally coordinate placental vasculogenesis. ^107–109^ These findings fit the paradigm that vascular abnormalities are driven by chronic hypoxia, oxidative stress, and dysregulated remodeling pathways.

### Placentas with both HCA and vascular abnormalities become more inflammatory

When HCA co-exists with vascular abnormalities, the Hallmark profile becomes more inflammatory. The 12 up-regulated sets—including Complement, Interferon α/γ, TNFα, ROS, Apoptosis, and Heme Metabolism—highlight a synergistic exacerbation of innate immunity, oxidative stress, and programmed cell death. ^110^ The exclusive down-regulation of DNA Repair suggests compromised genomic integrity, which can contribute to premature placental senescence and preterm labor. ^111, 112^ These data support the hypothesis that infectious triggers (HCA) amplify the detrimental vascular pathology, leading to a feed forward loop that accelerates parturition.

### Shared and opposing Hallmark pathways across phenotypes highlight distinct yet overlapping pathogenic mechanisms

The pathway signature of sPTB shares a core Hallmark pathway set with both infectious (HCA) and ischemic (vascular lesion) placental injury, yet the directionality of regulation diverges markedly, underscoring distinct mechanistic contributions of inflammation, oxidative stress, and cellular remodeling in each context. In both sPTB and HCA, KRAS signaling is down-regulated, a change that has been linked to impaired trophoblast proliferation and invasion, ^33, 113^ while the ROS and Interferon-γ pathways are inversely regulated—down-regulated in sPTB but up-regulated in HCA—suggesting that sterile inflammation in sPTB may rely less on classical oxidative and Th1-type antiviral responses and more on alternative, perhaps hypoxia-driven, inflammatory signals. ^12, 114, 115^ The shared up-regulation of epithelial-mesenchymal transition (EMT) in both sPTB and vascular abnormalities reflects a common dysregulation of trophoblast plasticity and extracellular matrix remodeling that can compromise placental barrier integrity. ^116^ The opposite regulation of protein secretion, oxidative phosphorylation, TGF-β signaling, allograft rejection, and mTORC1 signaling in sPTB versus vascular abnormalities underscores a shift from a catabolic, hypoxia-adapted state in sPTB to a pro-inflammatory, anabolic phenotype in vascular injury, mirroring the metabolic reprogramming in atherosclerotic vasculopathy. ^117^ When both HCA and vascular abnormalities coexist, the predominance of up-regulated EMT alongside down-regulation of interferon-α/β, complement, ROS, and allograft rejection pathways mirrors a compounded inflammatory milieu that may synergistically precipitate preterm labor. ^12, 118, 119^ Collectively, these findings highlight that while sPTB, HCA, and vascular abnormalities converge on key pathways of oxidative stress and immune modulation, the distinct directionality of their regulation reflects differential pathophysiological mechanisms that could inform targeted therapeutic strategies.

### Preterm placental circadian signatures

The functional pathway signatures uncovered in the pre-term placentas show a striking decoupling of the core circadian clock machinery from the downstream processes that govern the physiological “sleep-wake” cycle. In particular, the up-regulation of the GO:BP term Regulation of Circadian Sleep Wake Cycle indicates that genes normally involved in translating the central clock signal into organism-wide behavioral rhythms (for example, CIRBP, NR1D1, RORA, NPAS2, and NPAS3) are being transcriptionally enhanced. At the same time, the Reactome pathway Circadian Clock is down-regulated, implying that the core transcriptional–translation feedback loop comprising CLOCK, BMAL1 (ARNTL), PER1/2/3, and CRY1/2 is suppressed. Such a dissociation has been previously documented in other tissues from eight mice or humans exposed to chronic circadian mis-alignment, where clock output genes remain active while core clock components are dampened (Fagiani et al., 2022). ^120–123^ In the context of the placenta, this pattern suggests that the organ has partially retained its ability to coordinate “behavioral” or metabolic rhythms (e.g., nutrient transfer, hormone secretion) while its intrinsic time-keeping is compromised.

The down-regulated transcription factor target (TFT) gene sets associated with THRAP3, POU2AF1, ZNF146, and ARID5B further support the notion that the preterm placenta suffers from a loss of regulatory fidelity. THRAP3 (also known as TRAP220) is a co-activator of nuclear receptors that has been shown to modulate BMAL1–CLOCK activity and to integrate metabolic cues into circadian gene expression. ^124^ Other three transcription factor genes - POU2AF1, ZNF146, and ARID5B have been evidenced to be rhythmically expressed in blood of humans with insufficient sleep. ^59, 125^ The coordinated down-regulation of these TFTs indicates a failure to maintain the proper transcriptional network that sustains circadian oscillations. In preterm placentas this could manifest as aberrant timing of key events such as trophoblast invasion, nutrient transport, and placental hormone secretion—all processes that are known to be circadian-modulated. ^123, 126–128^

Clinically, disrupted circadian regulation in the placenta has been associated with a higher risk of preterm labor. Maternal circadian mis-alignment—whether through shift work, irregular sleep patterns, or jet lag—has been linked to increased levels of inflammatory cytokines and oxidative stress in the maternal–fetal unit, both of which are recognized precipitants of preterm birth. ^127, 129^ The data presented here suggest that the pre-term placenta may be both a sink for circadian mis-alignment (by down-regulating core clock components) and a source of maladaptive signaling (by up-regulating sleep-wake cycle regulators that are no longer properly timed). The loss of proper clock-gene regulation could also compromise the placenta’s ability to respond to the surge in cortisol that typically triggers parturition at term, thereby precipitating earlier labor. Further functional studies will be required to determine whether restoring normal circadian dynamics in the placenta can delay labor and improve neonatal outcomes.

### sPTB-enriched placental cell types and developmental pathways

The over-representation of trophoblast giant cells (TGCs) in spontaneous preterm birth (sPTB) samples is striking. TGCs are involved in implantation, hormone secretion, and vascular remodeling. ^130, 131^ Their enrichment suggests a compensatory response to the pronounced loss of extravillous trophoblasts (EVTs) that we also observed. EVTs are the primary mediators of arterial remodeling, and their depletion is consistently linked to inadequate uteroplacental perfusion and preterm labor. ^132, 133^ A shift toward the more primitive, highly proliferative TGC phenotype could therefore reflect an attempt to maintain vascular remodeling in the face of EVT dysfunction, yet may simultaneously disturb the delicate balance of invasion and differentiation that is essential for normal placental maturation. ^134, 135^

The IGFBP1- and DKK1-positive fetal placental cells that are up-regulated in sPTB add a second layer of functional disturbance. IGFBP1 is a secreted antagonist of insulin-like growth factor (IGF) signaling that modulates trophoblast proliferation, invasion, and hormone secretion. ^136^ Elevated IGFBP1 in maternal serum has been associated with sPTB and with a higher likelihood of cervical shortening. ^137^ DKK1 is an extracellular inhibitor of Wnt/β-catenin signaling. ^138^ Up-regulation of DKK1 in the fetal compartment could re-inhibit Wnt signaling, leading to impaired syncytiotrophoblast fusion and diminished production of progesterone and hCG—hormones that are crucial for maintaining uterine quiescence. ^139^ Thus, the simultaneous rise in IGFBP1 and DKK1 may converge on a common axis of reduced trophoblast differentiation and hormonal insufficiency, both of which are hallmarks of early parturition. ^140–142^

The down-regulation of syncytiotrophoblasts (STBs) and villous cytotrophoblasts (VCTs) is in line with previous transcriptomic studies that reported decreased trophoblast lineage signatures in preterm placentas. ^143–145^ STBs are the primary endocrine interface of the placenta; they produce and secrete hormones, vitamins, and growth factors essential for fetal growth and maternal adaptation. A loss of STB mass or function can lead to fetal growth restriction and a premature activation of the uterine contractile machinery. ^141, 146^ The concomitant reduction in VCTs, which serve as the proliferative reserve for STB formation, may exacerbate this effect by limiting the pool of cells available for renewal. ^147–149^

The depletion of fetal placental myeloid cells suggests a shift in the local immune environment. Placental macrophages (Hofbauer cells) play a pivotal role in trophoblast-myeloid crosstalk, modulating angiogenesis and immune tolerance. ^150–153^ Lower myeloid cell numbers may increase placental inflammation, ^154–156^ which has been recognized as a major precipitant of sPTB. ^77, 157^ It is possible that the observed myeloid reduction is a secondary consequence of the altered trophoblast landscape, further undermining the anti-inflammatory milieu required for a healthy pregnancy.

The gene-ontology terms related to maternal, placental, and embryonic development that are down-regulated point to a global developmental impairment. This aligns with earlier work demonstrating that dysregulated placental angiogenesis and impaired villous branching are frequent in preterm deliveries. ^158–162^ The convergence of decreased trophoblast differentiation, impaired hormone production, and disrupted angiogenic signaling creates a feed-forward loop that accelerates the transition from the proliferative growth phase to the functional maturation required for term labor.

### Correlations between sPTB-enriched Hallmark pathways and sPTB-enriched circadian pathways/TFT sets

The gene-sets that capture the canonical circadian clock as well as transcription factor targets of ARID5B, POU2AF1, and ZNF146 show robust correlations with a host of PTB-enriched Hallmark pathways that drive protein secretion, oxidative phosphorylation, androgen signaling, complement activation, ROS production, MYC-v1 transcription, TGF-β signaling, mTORC1 activity, and aberrant KRAS pathway activation. These pathways collectively represent a metabolic-inflammatory axis that has been linked to premature labor onset. ^163, 164^ Together, these results suggest that dysregulation of the circadian transcriptional network may amplify pro-inflammatory and oxidative stress responses while concurrently dampening oncogenic KRAS signaling, thereby tipping the balance toward a preterm phenotype. Overall, the data identify a network where placental circadian signaling and molecular networks converge to regulate metabolic and inflammatory processes that are critical for sustaining term pregnancy, and whose perturbation may precipitate PTB.

### Research Implications

The present study highlights a constellation of actionable pathways that could be targeted to avert spontaneous preterm birth. First, the profound loss of secretory and metabolic pathway signatures in sPTB suggests that supplementing metabolic cofactors, antioxidants, and regulating the TGF-β/mTORC1 axis may restore placental functional competence. Second, the marked oxidative-stress and TNFα/NF-κB activation in HCA point to a role for anti-inflammatory agents—TNFα/NF-κB inhibitors and reactive-oxygen-species scavengers—in tempering this surge. Third, disrupted placental circadian circuitry offers a rationale for chronotherapeutic strategies, such as timed light exposure or pharmacologic activators of core clock components (e.g., CRY or CLOCK modulators), to reestablish metabolic rhythms and delay premature uterine activation. Fourth, the cell-type shifts observed in sPTB—loss of extravillous and syncytiotrophoblasts—indicate that trophoblast-supportive growth factors or epigenetic modulators may preserve these essential lineages and prevent developmental arrest. A coordinated, multi-modal intervention program that couples serial placental transcriptomic monitoring with precision modulation of metabolism, inflammation, vascular remodeling, and circadian entrainment holds the most promise for reducing the incidence of spontaneous preterm birth.

## Strengths and Limitations

### Strengths

The POUCH study’s prospective design, comprehensive placental pathology assessment, and availability of FFPE specimens provide an unparalleled platform to perform high-resolution transcriptomics in a real-world population of spontaneous preterm birth (sPTB) cases and term controls. EdgeSeq™ technology, with its nuclease-protected probe-based amplification, allows for robust quantification of messenger RNA from FFPE tissues, thereby overcoming the limitations of RNA degradation inherent to paraffin fixation. ^165^ All samples were analyzed within a single batch and processed by a single laboratory. Concentrating on a demographically homogeneous subset of women aged 23–33 y increased internal validity. Rather than focusing on individual genes, we emphasized functional pathways. Use of the Hallmark gene sets and ranked log FCs of all genes for GSEA provides a more robust, noise-reduced enrichment analysis, avoiding arbitrary thresholds and redundancy of conventional DEGs. ^54, 56^ Curated placenta-specific circadian pathways, transcription-factor-target (TFT) gene sets, cell-type signatures, and developmental pathways further refine the analysis, enhancing tissue specificity and mechanistic interpretability. ^55, 59, 62^

### Limitations

The relatively small number of samples (N = 32) limits statistical power with the ability to detect subtle effects. Restricting the analysis to white women aged 23–33 y, while enhancing internal validity, reduces generalizability. Our future work intends to validate the findings in the larger and diverse sociodemographic context of the full POUCH cohort and to investigate systemic exposures that may contribute to the causality of the placental clock dysregulation.

## Conclusion

Spontaneous preterm birth is driven by a reciprocal interplay between chronic inflammation and hypoxic-ischemic placental programs, with distinct transcriptomic signatures in HCA versus vascular abnormalities as well as suppressed immune, perfusion and hormonal function and chrono-disturbance in the absence of major tissue damage. Disrupted circadian regulation and altered trophoblast subpopulations should be further studied to establish their connectivity and cell-type specific contribution to compromised placental function. These findings highlight mechanistic targets—metabolic, immune, and circadian pathways—and underscore the need for cell-type-specific diagnostics and chronotherapeutic interventions to reduce sPTB risk.

## Abbreviations

sPTB: spontaneous preterm birth
HCA: histologic chorioamnionitis
MV-O: maternal vascular obstructive
FV-O: fetal vascular obstructive
MV-I: maternal bleeding/vessel integrity
MV-D: lack of physiologic conversion of maternal spiral arteries
FV-I: fetal bleeding/vessel integrity
DV200: percentage of RNA fragments > 200 nt
GSEA: Gene Set Enrichment Analysis
DESeq2: differential expression analysis
HTG EdgeSeq: targeted transcriptome platform.

## References

1. Ananth CV, Vintzileos, A.M. Epidemiology of preterm birth and its clinical subtypes. J Matern Fetal Neonatal Med. 2006;19(12):773–782.

2. Wilcox AJ. Gestational age and preterm delivery. In Wilcox AJ, (Ed). Fertility and Pregnancy: An Epidemiologic Perspective: Oxford University Press 2010.

3. Manuck TA, Esplin MS, Biggio J, et al. The phenotype of spontaneous preterm birth: application of a clinical phenotyping tool. Am J Obstet Gynecol. 2015;212(4):487.e481–487.e411.

4. Goldenberg RL, Culhansswe JF, Iams JD, et al. Epidemiology and causes of preterm birth. The Lancet. 2008;371(9606):75–84.

5. Jesse DE, Swanson MS, Newton ER, et al. Racial Disparities in Biopsychosocial Factors and Spontaneous Preterm Birth Among Rural Low-Income Women. Journal of Midwifery & Women’s Health. 2009;54(1):35–42.

6. Salow AD, Pool LR, Grobman WA, et al. Associations of neighborhood-level racial residential segregation with adverse pregnancy outcomes. American Journal of Obstetrics & Gynecology. 2018;218(3):351.e351–351.e357.

7. Ohuma EO, Moller A-B, Bradley E, et al. National, regional, and global estimates of preterm birth in 2020, with trends from 2010: a systematic analysis. The Lancet. 2023;402(10409):1261–1271.

8. Brink LT, Roberts DJ, Wright CA, et al. Placental pathology in spontaneous and iatrogenic preterm birth: Different entities with unique pathologic features. Placenta. 2022;126:54–63.

9. Duan J, Xu F, Zhu C, et al. Histological chorioamnionitis and pathological stages on very preterm infant outcomes. Histopathology. 2024;84(6):1024–1037.

10. Hartnett ME. The effects of hypoxia, hyperoxia, and oxygen fluctuations on oxidative signaling in the preterm infant and on retinopathy of prematurity. Oxidative Stress and Antioxidant Protection 2016:77–92.

11. Menon R, Behnia F, Polettini J, et al. Novel pathways of inflammation in human fetal membranes associated with preterm birth and preterm pre-labor rupture of the membranes. Semin Immunopathol. 2020;42(4):431–450.

12. Areia AL, Mota-Pinto A. Inflammation and Preterm Birth: A Systematic Review. Reproductive Medicine. 2022;3(2):101–111.

13. Waring GJ, Robson SC, Bulmer JN, et al. Inflammatory Signalling in Fetal Membranes: Increased Expression Levels of TLR 1 in the Presence of Preterm Histological Chorioamnionitis. PLoS One. 2015;10(5):e0124298.

14. Leimert KB, Xu W, Princ MM, et al. Inflammatory Amplification: A Central Tenet of Uterine Transition for Labor. Front Cell Infect Microbiol. 2021;11:660983.

15. Kelly R, Holzman C, Senagore P, et al. Placental vascular pathology findings and pathways to preterm delivery. Am J Epidemiol. 2009;170(2):148–158.

16. Ciampa EJ, Flahardy P, Srinivasan H, et al. Hypoxia-inducible factor 1 signaling drives placental aging and can provoke preterm labor. Elife. 2023;12.

17. Li X, Liu X, Meng Q, et al. Circadian clock disruptions link oxidative stress and systemic inflammation to metabolic syndrome in obstructive sleep apnea patients. Front Physiol. 2022;13:932596.

18. Diallo AB, Coiffard B, Desbriere R, et al. Disruption of the Expression of the Placental Clock and Melatonin Genes in Preeclampsia. Int J Mol Sci. 2023;24(3).

19. Lin Y, He L, Cai Y, et al. The role of circadian clock in regulating cell functions: implications for diseases. MedComm. 2024;5(3):e504.

20. de Assis LVM, Kramer A. Circadian de(regulation) in physiology: implications for disease and treatment. Genes Dev. 2024;38(21-24):933–951.

21. Khan S, Duan P, Yao L, et al. Shiftwork-Mediated Disruptions of Circadian Rhythms and Sleep Homeostasis Cause Serious Health Problems. International Journal of Genomics. 2018;2018(1):8576890.

22. Fishbein AB, Knutson KL, Zee PC. Circadian disruption and human health. The Journal of Clinical Investigation. 2021;131(19).

23. Moškon M, Kovač U, Raspor Dall’Olio L, et al. Circadian characteristics of term and preterm labors. Sci Rep. 2024;14(1):4033.

24. Suzumori N, Ebara T, Matsuki T, et al. Effects of long working hours and shift work during pregnancy on obstetric and perinatal outcomes: A large prospective cohort study—Japan Environment and Children’s Study. Birth. 2020;47(1):67–79.

25. Nikhil KL, Bates K, Sapiro E, et al. Fetoplacental Circadian Rhythms Develop and Then Synchronize to the Mother In Utero. J Biol Rhythms. 2026:7487304261435435.

26. Ahmadi SM, Perez ML, Guardia CM. Secretion of placental peptide hormones: functions and trafficking. Frontiers in Endocrinology. 2025;Volume 16 - 2025.

27. Basak S, Varma S, Duttaroy A. Modulation of fetoplacental growth, development and reproductive function by endocrine disrupters. Frontiers in Endocrinology. 2023;14.

28. Duong TV, Yaw AM, Zhou G, et al. Interaction between time-of-day and oxytocin efficacy in mice and humans with and without gestational diabetes. bioRxiv. 2024.

29. Li H, Peng H, Hong W, et al. Human Placental Endothelial Cell and Trophoblast Heterogeneity and Differentiation Revealed by Single-Cell RNA Sequencing. Cells. 2023;12(1):87.

30. Liu Y, Fan X, Wang R, et al. Single-cell RNA-seq reveals the diversity of trophoblast subtypes and patterns of differentiation in the human placenta Cell Research. 2018;28(8):819–832.

31. Derisoud E, Jiang H, Zhao A, et al. Revealing the molecular landscape of human placenta: a systematic review and meta-analysis of single-cell RNA sequencing studies. Hum Reprod Update. 2024;30(4):410–441.

32. Dietrich B, Haider S, Meinhardt G, et al. WNT and NOTCH signaling in human trophoblast development and differentiation. Cell Mol Life Sci. 2022;79(6):292.

33. Liu L, Tang L, Chen S, et al. Decoding the molecular pathways governing trophoblast migration and placental development; a literature review. Frontiers in Endocrinology. 2024;Volume 15 - 2024.

34. Giaglis S, Stoikou M, Grimolizzi F, et al. Neutrophil migration into the placenta: Good, bad or deadly? Cell Adh Migr. 2016;10(1-2):208–225.

35. Lv M, Jia Y, Dong J, et al. The landscape of decidual immune cells at the maternal–fetal interface in parturition and preterm birth. Inflammation Research. 2025;74(1):44.

36. Alippe Y, Hatterschide J, Coyne CB, et al. Innate immune responses to pathogens at the maternal–fetal interface. Nature Reviews Immunology. 2025;25(12):869–884.

37. Holzman C, Bullen B, Fisher R, et al. Pregnancy outcomes and community health: the POUCH study of preterm birth. Paediatric and Perinatal Epidemiology. 2001;15(s2):136–158.

38. Holzman C, Senagore PK, Wang J. Mononuclear leukocyte infiltrate in extraplacental membranes and preterm delivery. Am J Epidemiol. 2013;177(10):1053–1064.

39. Holzman C, Lin X, Senagore P, et al. Histologic Chorioamnionitis and Preterm Delivery. American Journal of Epidemiology. 2007;166(7):786–794.

40. Gargano JW, Holzman C, Senagore P, et al. Mid-pregnancy circulating cytokine levels, histologic chorioamnionitis and spontaneous preterm birth. J Reprod Immunol. 2008;79(1):100–110.

41. Borchert S, Herold T, Kalbourtzis S, et al. Transcriptome-Wide Gene Expression Profiles from FFPE Materials Based on a Nuclease Protection Assay Reveals Significantly Different Patterns between Synovial Sarcomas and Morphologic Mimickers. Cancers. 2022;14(19):4737.

42. Koll FJ, Metzger E, Hamann J, et al. Overexpression of KMT9α Is Associated with Aggressive Basal-like Muscle-Invasive Bladder Cancer. Cells. 2023;12(4).

43. Koll FJ, Döring C, Olah C, et al. Optimizing identification of consensus molecular subtypes in muscle-invasive bladder cancer: a comparison of two sequencing methods and gene sets using FFPE specimens. BMC Cancer. 2023;23(1):504.

44. Díez-Ahijado L, del Rincón AM, Marimón L, et al. HTGAnalyzer: An accessible R package with a web interface for enhanced transcriptomic analysis in precision medicine. Computers in Biology and Medicine. 2025;196:110772.

45. Fernández-Serra A, López-Reig R, Romero I, et al. Comparative evaluation of HTG and TempO Seq targeted transcriptome profiling methods. Scientific Reports. 2026;16(1):6108.

46. Bankhead P, Loughrey MB, Fernández JA, et al. QuPath: Open source software for digital pathology image analysis. Scientific Reports. 2017;7(1):16878.

47. Love MI, Huber W, Anders S. Moderated estimation of fold change and dispersion for RNA-seq data with DESeq2. Genome Biology. 2014;15(12):550.

48. Ritchie ME, Phipson B, Wu D, et al. limma powers differential expression analyses for RNA-sequencing and microarray studies. Nucleic Acids Research. 2015;43(7):e47–e47.

49. Sood R, Zehnder JL, Druzin ML, et al. Gene expression patterns in human placenta. Proceedings of the National Academy of Sciences. 2006;103(14):5478–5483.

50. Pique-Regi R, Romero R, Tarca AL, et al. Single cell transcriptional signatures of the human placenta in term and preterm parturition. eLife. 2019;8:e52004.

51. Paquette AG, MacDonald J, Bammler T, et al. Placental transcriptomic signatures of spontaneous preterm birth. Am J Obstet Gynecol. 2023;228(1):73.e71–73.e18.

52. Schoenmakers S, Aagaard K, Borenstein-Levin L, et al. Editorial: Preterm birth and placental pathology. Front Endocrinol (Lausanne). 2023;14:1168185.

53. Smyth GK. limma: Linear Models for Microarray Data. In Gentleman R. CV, Dudoit S., Irizarry R., Huber W., (Ed). Bioinformatics and Computational Biology Solutions Using R and Bioconductor Statistics for Biology and Health. New York: Springer 2005.

54. Subramanian A, Tamayo P, Mootha VK, et al. Gene set enrichment analysis: A knowledge-based approach for interpreting genome-wide expression profiles. Proceedings of the National Academy of Sciences. 2005;102(43):15545–15550.

55. Castanza AS, Recla JM, Eby D, et al. Extending support for mouse data in the Molecular Signatures Database (MSigDB). Nature Methods. 2023;20(11):1619–1620.

56. Liberzon A, Birger C, Thorvaldsdóttir H, et al. The Molecular Signatures Database (MSigDB) hallmark gene set collection. Cell Syst. 2015;1(6):417–425.

57. Dolgalev I. msigdbr: MSigDB Gene Sets for Multiple Organisms in a Tidy Data Format. R package version 2610 2026.

58. Yu G, Wang L-G, Han Y, et al. clusterProfiler: an R package for comparing biological themes among gene clusters. OMICS. 2012;16(5):284–287.

59. Li S, Shui K, Zhang Y, et al. CGDB: a database of circadian genes in eukaryotes. Nucleic Acids Research. 2017;45(D1):D397–D403.

60. Hänzelmann S, Castelo R, Guinney J. GSVA: gene set variation analysis for microarray and RNA-Seq data. BMC Bioinformatics. 2013;14(1):7.

61. Harrell FE. Hmisc: Harrell Miscellaneous. R package version 52–5 2003.

62. Wang T, Roach MJ, Harvey K, et al. snPATHO-seq, a versatile FFPE single-nucleus RNA sequencing method to unlock pathology archives. Communications Biology. 2024;7(1):1340.

63. Sarantopoulou D, Tang SY, Ricciotti E, et al. Comparative evaluation of RNA-Seq library preparation methods for strand-specificity and low input. Sci Rep. 2019;9(1):13477.

64. Illumina. Guidelines for obtaining high-quality RNA sequencing results from degraded RNA with Illumina RNA enrichment assays 2016.

65. Elshenawy S, Pinney SE, Stuart T, et al. The Metabolomic Signature of the Placenta in Spontaneous Preterm Birth. Int J Mol Sci. 2020;21(3).

66. Lien YC, Zhang Z, Cheng Y, et al. Human Placental Transcriptome Reveals Critical Alterations in Inflammation and Energy Metabolism with Fetal Sex Differences in Spontaneous Preterm Birth. Int J Mol Sci. 2021;22(15).

67. Voros C, Stavros S, Sapantzoglou I, et al. The Role of Placental Mitochondrial Dysfunction in Adverse Perinatal Outcomes: A Systematic Review. J Clin Med. 2025;14(11).

68. Cesana M, Tufano G, Panariello F, et al. TFEB controls syncytiotrophoblast formation and hormone production in placenta. Cell Death & Differentiation. 2024;31(11):1439–1451.

69. Sarig G, Brenner B. Coagulation, inflammation, and pregnancy complications. The Lancet. 2004;363(9403):96–97.

70. Heuser CC BD. Disorders of Coagulation in Pregnancy. In James D SP, Weiner C, Gonik B, Robson S, (Ed). High-Risk Pregnancy: Management Options: Five-Year Institutional Subscription with Online Updates: Cambridge University Press 2017:1085–1107.

71. Kumar V, Stewart JH. The complement system in human pregnancy and preeclampsia. Frontiers in Immunology. 2025;Volume 16 - 2025.

72. Lin Z, Wu S, Jiang Y, et al. Unraveling the molecular mechanisms driving enhanced invasion capability of extravillous trophoblast cells: a comprehensive review. Journal of Assisted Reproduction and Genetics. 2024;41(3):591–608.

73. Vornic I, Caprariu R, Novacescu D, et al. Molecular Insights into Human Placentation: From Villous Morphogenesis to Pathological Pathways and Translational Biomarkers. International Journal of Molecular Sciences. 2025;26(19):9483.

74. Ma J, Gong F, Kim E, et al. Early elevations of RAS protein level and activity are critical for the development of PDAC in the context of inflammation. Cancer Lett. 2024;586:216694.

75. Singh N, Herbert B, Sooranna G, et al. Distinct preterm labor phenotypes have unique inflammatory signatures and contraction associated protein profiles†. Biol Reprod. 2019;101(5):1031–1045.

76. Fasoulakis Z, Koutras A, Ntounis T, et al. Inflammatory Molecules Responsible for Length Shortening and Preterm Birth. Cells. 2023;12(2).

77. Georges HM, Norwitz ER, Abrahams VM. Predictors of Inflammation-Mediated Preterm Birth. Physiology. 2025;40(1):26–36.

78. Oghbaei F, Zarezadeh R, Jafari-Gharabaghlou D, et al. Epithelial-mesenchymal transition process during embryo implantation. Cell and Tissue Research. 2022;388(1):1–17.

79. de Castro Silva M, Richardson LS, Kechichian T, et al. Inflammation, but not infection, induces EMT in human amnion epithelial cells. Reproduction. 2020;160(4):627–638.

80. Menon R. Epithelial to mesenchymal transition (EMT) of feto-maternal reproductive tissues generates inflammation: a detrimental factor for preterm birth. BMB Rep. 2022;55(8):370–379.

81. Ackerman IV WE, Rigo MM, DaSilva-Arnold SC, et al. Epigenetic Changes Regulating Epithelial–Mesenchymal Plasticity in Human Trophoblast Differentiation. Cells. 2025;14(13):970.

82. Jang D-i, Lee A-H, Shin H-Y, et al. The Role of Tumor Necrosis Factor Alpha (TNF-α) in Autoimmune Disease and Current TNF-α Inhibitors in Therapeutics. International Journal of Molecular Sciences. 2021;22(5):2719.

83. Martin LF MN, Polettini J, et al. Histologic Chorioamnionitis Induces Higher Lipid Oxidative Damage in Amniochorion Membranes from Pregnancies Complicated by Spontaneous Prematurity. Reproductive Sciences. 2017;24:105A–105A.

84. Aouache R, Biquard L, Vaiman D, et al. Oxidative Stress in Preeclampsia and Placental Diseases. International Journal of Molecular Sciences. 2018;19(5):1496.

85. Sultana Z, Qiao Y, Maiti K, et al. Involvement of oxidative stress in placental dysfunction, the pathophysiology of fetal death and pregnancy disorders. Reproduction. 2023;166(2):R25–R38.

86. Budal EB, Bentsen MHL, Kessler J, et al. Histologic chorioamnionitis in extremely preterm births, microbiological findings and infant outcome. The Journal of Maternal-Fetal & Neonatal Medicine. 2023;36(1):2196599.

87. Kim YM, Romero R, Chaiworapongsa T, et al. Toll-like receptor-2 and −4 in the chorioamniotic membranes in spontaneous labor at term and in preterm parturition that are associated with chorioamnionitis. American Journal of Obstetrics and Gynecology. 2004;191(4):1346–1355.

88. Firmal P, Shah VK, Chattopadhyay S. Insight Into TLR4-Mediated Immunomodulation in Normal Pregnancy and Related Disorders. Front Immunol. 2020;11:807.

89. Dash S, Sahu AK, Srivastava A, et al. Exploring the extensive crosstalk between the antagonistic cytokines- TGF-β and TNF-α in regulating cancer pathogenesis. Cytokine. 2021;138:155348.

90. Massagué J, Sheppard D. TGF-&#x3b2; signaling in health and disease. Cell. 2023;186(19):4007–4037.

91. Zhang M, Liu Q, Meng H, et al. Ischemia-reperfusion injury: molecular mechanisms and therapeutic targets. Signal Transduction and Targeted Therapy. 2024;9(1):12.

92. Horvat Mercnik M, Schliefsteiner C, Sanchez-Duffhues G, et al. TGFβ signalling: a nexus between inflammation, placental health and preeclampsia throughout pregnancy. Hum Reprod Update. 2024;30(4):442–471.

93. Shakir D, Batie M, Kwok CS, et al. NF-κB is a central regulator of hypoxia-induced gene expression. EMBO Rep. 2026;27(2):416–432.

94. Tao Y, Zhang M, Chen L, et al. Macrophage AMPK activated by oxidative stress drives profibrotic crosstalk with tubular cells to accelerate renal fibrosis after ischemic and reperfusion injury. Redox Biology. 2026;90:104002.

95. Yi SG, Gaber AO, Chen W. B-cell response in solid organ transplantation. Frontiers in Immunology. 2022;Volume 13 - 2022.

96. Duneton C, Winterberg PD, Ford ML. Activation and regulation of alloreactive T cell immunity in solid organ transplantation. Nature Reviews Nephrology. 2022;18(10):663–676.

97. Bournique E, Sanchez A, Oh S, et al. ATM and IRAK1 orchestrate two distinct mechanisms of NF-κB activation in response to DNA damage. Nature Structural & Molecular Biology. 2025;32(4):740–755.

98. (NTP) NTP. Hallmark Gene Sets. Chemical Effects in Biological Systems (CEBS). Research Triangle Park, NC (USA) 2026.

99. Meng LB, Chen K, Zhang YM, et al. Common Injuries and Repair Mechanisms in the Endothelial Lining. Chin Med J (Engl). 2018;131(19):2338–2345.

100. Barbosa GSB, Câmara NOS, Ledesma FL, et al. Vascular injury in glomerulopathies: the role of the endothelium. Front Nephrol. 2024;4:1396588.

101. Davies JE PJ, Yong HEJ, et al.. Epithelial-mesenchymal transition during extravillous trophoblast differentiation. CELL ADHESION & MIGRATION. 2016;10(3):310–321.

102. Piera-Velazquez S, Jimenez SA. Endothelial to Mesenchymal Transition: Role in Physiology and in the Pathogenesis of Human Diseases. Physiol Rev. 2019;99(2):1281–1324.

103. Choudhury J, Pandey D, Chaturvedi PK, et al. Epigenetic regulation of epithelial to mesenchymal transition: a trophoblast perspective. Molecular Human Reproduction. 2022;28(5).

104. Choudhury J, Dhole B, Aggarwal K, et al. Hypoxia regulates epithelial to mesenchymal transition-associated genes in human trophoblast cells by modulating DNA methylation. PLoS One. 2026;21(4):e0325053.

105. Kim R, Chang W. Endothelial-to-Mesenchymal Transition in Health and Disease: Molecular Insights and Therapeutic Implications. Int J Mol Sci. 2025;26(23).

106. Ali A, Yun S. Multifaceted Role of Notch Signaling in Vascular Health and Diseases. Biomedicines. 2025;13(4):837.

107. Holderfield MT, Hughes CC. Crosstalk between vascular endothelial growth factor, notch, and transforming growth factor-beta in vascular morphogenesis. Circ Res. 2008;102(6):637–652.

108. Zhao WX, Lin JH. Notch signaling pathway and human placenta. Int J Med Sci. 2012;9(6):447–452.

109. Dudley AC, Griffioen AW. Pathological angiogenesis: mechanisms and therapeutic strategies. Angiogenesis. 2023;26(3):313–347.

110. Janžič L, Kouter K. Complement and inflammasome crosstalk in chronic inflammation. Frontiers in Immunology. 2026;Volume 17 - 2026.

111. Menon R, Boldogh I, Urrabaz-Garza R, et al. Senescence of primary amniotic cells via oxidative DNA damage. PLoS One. 2013;8(12):e83416.

112. Singh VP, Singh P. Linking DNA damage and senescence to gestation period and lifespan in placental mammals. Frontiers in Cell and Developmental Biology. 2024;Volume 12 - 2024.

113. Carvajal L, Gutiérrez J, Morselli E, et al. Autophagy Process in Trophoblast Cells Invasion and Differentiation: Similitude and Differences With Cancer Cells. Front Oncol. 2021;11:637594.

114. Motomura K, Romero R, Plazyo O, et al. The alarmin S100A12 causes sterile inflammation of the human chorioamniotic membranes as well as preterm birth and neonatal mortality in mice†. Biol Reprod. 2021;105(6):1494–1509.

115. Baker BC, Heazell AEP, Sibley C, et al. Hypoxia and oxidative stress induce sterile placental inflammation in vitro. Sci Rep. 2021;11(1):7281.

116. Ma Z, Sagrillo-Fagundes L, Mok S, et al. Mechanobiological regulation of placental trophoblast fusion and function through extracellular matrix rigidity. Scientific Reports. 2020;10(1):5837.

117. Liu YX, Guo FM, Qiu WJ, et al. Metabolic Reprogramming and Cell Interaction in Atherosclerosis: From Molecular Mechanisms to Therapeutic Strategies. J Cardiovasc Dev Dis. 2025;12(10).

118. Green ES, Arck PC. Pathogenesis of preterm birth: bidirectional inflammation in mother and fetus. Semin Immunopathol. 2020;42(4):413–429.

119. Wu Y, Pei C, Huang J. The multifactorial and complex nature of preterm birth. The Journal of Maternal-Fetal & Neonatal Medicine. 2025;38.

120. Wolff CA, Gutierrez-Monreal MA, Meng L, et al. Defining the age-dependent and tissue-specific circadian transcriptome in male mice. Cell Reports. 2023;42(1).

121. Seya T, Koike N, Okubo N, et al. Chronic circadian misalignment accelerates sarcopenia progression in mice. Front Physiol. 2025;16:1686942.

122. Sutton E, Pekovic-Vaughan V. Time to Reset: The Interplay Between Circadian Rhythms and Redox Homeostasis in Skeletal Muscle Ageing and Systemic Health. Antioxidants. 2025;14(9):1132.

123. Fagiani F, Di Marino D, Romagnoli A, et al. Molecular regulations of circadian rhythm and implications for physiology and diseases. Signal Transduct Target Ther. 2022;7(1):41.

124. Jang H-J, Lee YH, Dao T, et al. Thrap3 promotes nonalcoholic fatty liver disease by suppressing AMPK-mediated autophagy. Experimental & Molecular Medicine. 2023;55(8):1720–1733.

125. Möller-Levet CS, Archer SN, Bucca G, et al. Effects of insufficient sleep on circadian rhythmicity and expression amplitude of the human blood transcriptome. Proceedings of the National Academy of Sciences. 2013;110(12):E1132–E1141.

126. Reiter RJ, Tan DX, Korkmaz A, et al. Melatonin and stable circadian rhythms optimize maternal, placental and fetal physiology. Human Reproduction Update. 2014;20(2):293–307.

127. Yao N, Kinouchi K, Katoh M, et al. Maternal circadian rhythms during pregnancy dictate metabolic plasticity in offspring. Cell Metabolism. 2025;37(2):395–412.e396.

128. Gao Y, Yu X, Wang Y, et al. Metabolic control of feto–placental development and pregnancy outcomes. Nature Reviews Endocrinology. 2026;22(3):153–165.

129. Cui Z, Xu H, Wu F, et al. Maternal circadian rhythm disruption affects neonatal inflammation via metabolic reprograming of myeloid cells. Nature Metabolism. 2024;6(5):899–913.

130. Hu D, Cross JC. Development and function of trophoblast giant cells in the rodent placenta. Int J Dev Biol. 2010;54(2-3):341–354.

131. Favaron PO, Carter AM. The trophoblast giant cells of cricetid rodents. Frontiers in Cell and Developmental Biology. 2023;Volume 10 - 2022.

132. Pollheimer J, Vondra S, Baltayeva J, et al. Regulation of Placental Extravillous Trophoblasts by the Maternal Uterine Environment. Front Immunol. 2018;9:2597.

133. Wei X-W, Zhang Y-C, Wu F, et al. The role of extravillous trophoblasts and uterine NK cells in vascular remodeling during pregnancy. Frontiers in Immunology. 2022;Volume 13 - 2022.

134. Huppertz B. Placental Development with Histological Aspects. In Huppertz B, Schleußner, E., (Ed). The Placenta. Berlin, Heidelberg: Berlin, Heidelberg 2023.

135. Vornic I, Buciu V, Furau CG, et al. The Interplay of Molecular Factors and Morphology in Human Placental Development and Implantation. Biomedicines. 2024;12(12):2908.

136. Li X, Li C, Wang Y, et al. IGFBP1 inhibits the invasion, migration, and apoptosis of HTR-8/SVneo trophoblast cells in preeclampsia. Hypertension in Pregnancy. 2022;41(1):53–63.

137. Larisa Mešić Ð, Dragana M, Feđa O, et al. IGFBP-1 marker of cervical ripening and predictor of preterm birth. Medicinski glasnik. 2016;13(2).

138. Jiang H, Zhang Z, Yu Y, et al. Drug Discovery of DKK1 Inhibitors. Frontiers in Pharmacology. 2022;Volume 13 - 2022.

139. Kim M, Jang YJ, Lee M, et al. The transcriptional regulatory network modulating human trophoblast stem cells to extravillous trophoblast differentiation. Nature Communications. 2024;15(1):1285.

140. Handwerger S. New insights into the regulation of human cytotrophoblast cell differentiation. Mol Cell Endocrinol. 2010;323(1):94–104.

141. Zhou H, Zhao C, Wang P, et al. Regulators involved in trophoblast syncytialization in the placenta of intrauterine growth restriction. Front Endocrinol (Lausanne). 2023;14:1107182.

142. Lawless L, Qin Y, Xie L, et al. Trophoblast Differentiation: Mechanisms and Implications for Pregnancy Complications. Nutrients. 2023;15(16):3564.

143. Szilagyi A, Gelencser Z, Romero R, et al. Placenta-Specific Genes, Their Regulation During Villous Trophoblast Differentiation and Dysregulation in Preterm Preeclampsia. Int J Mol Sci. 2020;21(2).

144. Akram KM, Kulkarni NS, Brook A, et al. Transcriptomic analysis of the human placenta reveals trophoblast dysfunction and augmented Wnt signalling associated with spontaneous preterm birth. Front Cell Dev Biol. 2022;10:987740.

145. Uhm C, Gu J, Ju W, et al. Single-nucleus RNA sequencing reveals distinct pathophysiological trophoblast signatures in spontaneous preterm birth subtypes. Cell & Bioscience. 2025;15(1):1.

146. Bi Y, Yang J, Li X, et al. Single-cell insights into trophoblast heterogeneity and adaptive dysfunction in selective fetal growth restriction. Communications Biology. 2026;9(1):387.

147. Aplin JD, Jones CJP. Cell dynamics in human villous trophoblast. Human Reproduction Update. 2021;27(5):904–922.

148. Renaud SJ, Jeyarajah MJ. How trophoblasts fuse: an in-depth look into placental syncytiotrophoblast formation. Cell Mol Life Sci. 2022;79(8):433.

149. Podinić T, MacAndrew, A., Raha, S.. Trophoblast Syncytialization: A Metabolic Crossroads. In Kloc M, Uosef, A., (Ed). Syncytia: Origin, Structure, and Functions Results and Problems in Cell Differentiation: Springer, Cham 2024.

150. Zulu Michael Z, Martinez Fernando O, Gordon S, et al. The Elusive Role of Placental Macrophages: The Hofbauer Cell. Journal of Innate Immunity. 2019;11(6):447–456.

151. Thomas JR, Appios A, Zhao X, et al. Phenotypic and functional characterization of first-trimester human placental macrophages, Hofbauer cells. Journal of Experimental Medicine. 2020;218(1).

152. Yang SW, Hwang HS, Kang YS. The role of placenta Hofbauer cells during pregnancy and pregnancy complications. Obstet Gynecol Sci. 2025;68(1):9–17.

153. Baráth BR, Bojcsuk D, Bene K, et al. Distinct transcriptional and epigenomic programs define Hofbauer cells in term placenta. JCI Insight. 2025;11(3).

154. Tami M, Hontecillas-Prieto L, García-Domínguez D, et al. Decreased number of myeloid-derived suppressor cells in the placental trophoblast of gestational diabetes mellitus. Possible role of leptin. Immunobiology. 2025;230(3):152897.

155. Pang B, Hu C, Li H, et al. Myeloidderived suppressor cells: Escorts at the maternal-fetal interface. Front Immunol. 2023;14:1080391.

156. Costa ML, Robinette ML, Bugatti M, et al. Two Distinct Myeloid Subsets at the Term Human Fetal-Maternal Interface. Front Immunol. 2017;8:1357.

157. Cervantes EM, Girard S. Placental Inflammation in Preterm Premature Rupture of Membranes and Risk of Neurodevelopmental Disorders. Cells. 2025;14(13):965.

158. Sakdapreecha L, Koonmee S, Triamwittayanon T, et al. Accelerated villous maturation of placentas in spontaneous preterm birth. Journal of the Medical Association of Thailand. 2017;100:1145–1149.

159. Huang Z, Huang S, Song T, et al. Placental Angiogenesis in Mammals: A Review of the Regulatory Effects of Signaling Pathways and Functional Nutrients. Advances in Nutrition. 2021;12(6):2415–2434.

160. Jaiman S, Romero R, Pacora P, et al. Disorders of placental villous maturation are present in one-third of cases with spontaneous preterm labor. J Perinat Med. 2021;49(4):412–430.

161. Jaiman S, Romero R, Bhatti G, et al. The role of the placenta in spontaneous preterm labor and delivery with intact membranes. J Perinat Med. 2022;50(5):553–566.

162. Jacobs A, Al-Juboori SI, Dobrinskikh E, et al. Placental differences between severe fetal growth restriction and hypertensive disorders of pregnancy requiring early preterm delivery: morphometric analysis of the villous tree supported by artificial intelligence. Am J Obstet Gynecol. 2024;231(5):552.e551–552.e513.

163. Yan Y, Gu Z, Li B, et al. Metabonomics profile analysis in inflammation-induced preterm birth and the potential role of metabolites in regulating premature cervical ripening. Reproductive Biology and Endocrinology. 2022;20(1):135.

164. Gomez-Lopez N, Galaz J, Miller D, et al. The immunobiology of preterm labor and birth: intra-amniotic inflammation or breakdown of maternal-fetal homeostasis. Reproduction. 2022;164(2):R11–r45.

165. Qi Z, Wang L, Desai K, et al. Reliable Gene Expression Profiling from Small and Hematoxylin and Eosin–Stained Clinical Formalin-Fixed, Paraffin-Embedded Specimens Using the HTG EdgeSeq Platform. The Journal of Molecular Diagnostics. 2019;21(5):796–807.

